# SARS-CoV-2 infection severity is linked to superior humoral immunity against the spike

**DOI:** 10.1101/2020.09.12.294066

**Authors:** Jenna J. Guthmiller, Olivia Stovicek, Jiaolong Wang, Siriruk Changrob, Lei Li, Peter Halfmann, Nai-Ying Zheng, Henry Utset, Christopher T. Stamper, Haley L. Dugan, William D. Miller, Min Huang, Ya-Nan Dai, Christopher A. Nelson, Paige D. Hall, Maud Jansen, Kumaran Shanmugarajah, Jessica S. Donington, Florian Krammer, Daved H. Fremont, Andrzej Joachimiak, Yoshihiro Kawaoka, Vera Tesic, Maria Lucia Madariaga, Patrick C. Wilson

**Affiliations:** Department of Medicine, Section of Rheumatology, University of Chicago, Chicago, IL 60637, USA; Influenza Research Institute, Department of Pathobiological Sciences, School of Veterinary Medicine, University of Wisconsin-Madison, Madison, WI, 53711; Committee on Immunology, University of Chicago, Chicago, IL 60637, USA; Department of Medicine, Section of Pulmonary and Critical Care Medicine, University of Chicago, Chicago, IL 60637, USA; Department of Pathology and Immunology and Center for Structural Genomics of Infectious Diseases, Consortium for Advanced Science and Engineering, Washington University School of Medicine, St. Louis, MO 63110, USA; Department of Medicine, University of Chicago, Chicago, IL 60637, USA; Department of Surgery, University of Chicago, Chicago, IL 60637, USA; Department of Microbiology, Icahn School of Medicine at Mount Sinai, New York, NY 10029, USA; Center for Structural Genomics of Infectious Diseases, Consortium for Advanced Science and Engineering, University of Chicago, Chicago, IL 60667; Structural Biology Center, X-ray Science Division, Argonne National Laboratory, Argonne, IL 60439, USA; Department of Pathology, University of Chicago, Chicago, IL 60637, USA

**Author notes:** These authors contributed equally (Jenna J. Guthmiller, Olivia Stovicek, Jiaolong Wang).

## Abstract

Severe acute respiratory syndrome coronavirus 2 (SARS-CoV-2) is currently causing a global pandemic. The antigen specificity and kinetics of the antibody response mounted against this novel virus are not understood in detail. Here, we report that subjects with a more severe SARS-CoV-2 infection exhibit a larger antibody response against the spike and nucleocapsid protein and epitope spreading to subdominant viral antigens, such as open reading frame 8 and non-structural proteins. Subjects with a greater antibody response mounted a larger memory B cell response against the spike, but not the nucleocapsid protein. Additionally, we revealed that antibodies against the spike are still capable of binding the D614G spike mutant and cross-react with the SARS-CoV-1 receptor binding domain. Together, this study reveals that subjects with a more severe SARS-CoV-2 infection exhibit a greater overall antibody response to the spike and nucleocapsid protein and a larger memory B cell response against the spike.

---

Entry of SARS-CoV-2 into host cells is mediated by surface trimeric spike protein via interaction between the spike receptor-binding domain (RBD) and angiotensin-converting enzyme 2^1,2^. SARS-CoV-2 expresses numerous potential antigens, including four structural proteins (spike, nucleocapsid (N) protein, matrix and envelope protein), 16 nonstructural proteins/antigens (NSP1–NSP16), and several accessory open reading frame (ORF) proteins, including ORF7 and ORF8^3,4^. To date, little is known about the specificities and kinetics of antibodies elicited in response to this infection and how coronavirus disease 2019 (COVID-19) severity relates to magnitude of the humoral immune response.

To address this critically important knowledge gap, we collected plasma samples from 35 hospitalized acutely SARS-CoV-2 infected subjects and 105 convalescent subjects^5^ (Supplemental Tables 1 and 2). Plasma was tested against the spike, N protein, ORF7a, ORF8, and NSP3, NSP9, NSP10, and NSP15 of SARS-CoV-2. Notably, all subjects within the acutely infected cohort were hospitalized, whereas only 9% (8/105) of subjects in the convalescent cohort had been hospitalized (Supplemental Tables 1 and 2). 89% of acutely infected subjects and 98% of convalescent subjects had detectable antibodies against one or more SARS-CoV-2 antigens (Fig. 1a), with nearly all subjects mounting a response against the spike and N protein (Fig. 1b). We further identified that convalescent subjects mounted a predominant response against the RBD of the spike protein and the RNA-binding domain of the N protein (Extended data Fig. 1a and b), suggesting these domains contain the immunodominant epitopes of these antigens. A larger frequency of acutely infected subjects mounted antibodies against ORF7a, ORF8, and NSP antigens (Fig. 1b, Extended data Fig. 1c) suggesting antibodies against these antigens are either short-lived or only induced by more severe infection. Moreover, the anti-N protein antibody response preceded the antibody response against the spike protein and was consistently higher across all time points, peaking between the 2^nd^ and 3^rd^ week after the onset of symptoms and retracting between the 3^rd^ and 4^th^ weeks after symptom onset (Fig. 1c and Extended data Fig. 1d). Furthermore, we found a strong positive correlation between the anti-N protein and anti-spike IgG titers in both the acutely infected and convalescent cohorts (Extended data 1e and f), indicating subjects who generally mounted a robust antibody response upon SARS-CoV-2 infection tended to mount a robust response against both antigens. Titers against both the spike and N protein persisted even 2+ months after symptom onset (Fig. 1c), indicating the antibody response against these two antigens is stable amongst subjects with symptomatic infection, a finding consistent with other reports^6,7^. We did not observe a statistical difference in antibody titers against the spike and N protein by individual subjects in either the acute or convalescent subject cohorts (Fig. 1d and e), likely due to dramatic subject-to-subject variation. However, antibody titers against the spike and N protein were significantly higher than antibody titers against ORF7a and ORF8 (Fig. 1d and e). Together, these data reveal the antibody response against SARS-CoV-2 is largely driven against the spike and N protein and that the anti-N protein antibody response precedes the response against the spike. The differences in the kinetics and magnitude of the N protein response are potentially due to the differences in protein expression, as N protein completely covers the entire viral genome, whereas a single virion only expresses ∼26 trimers^8^.

**Fig. 1:**
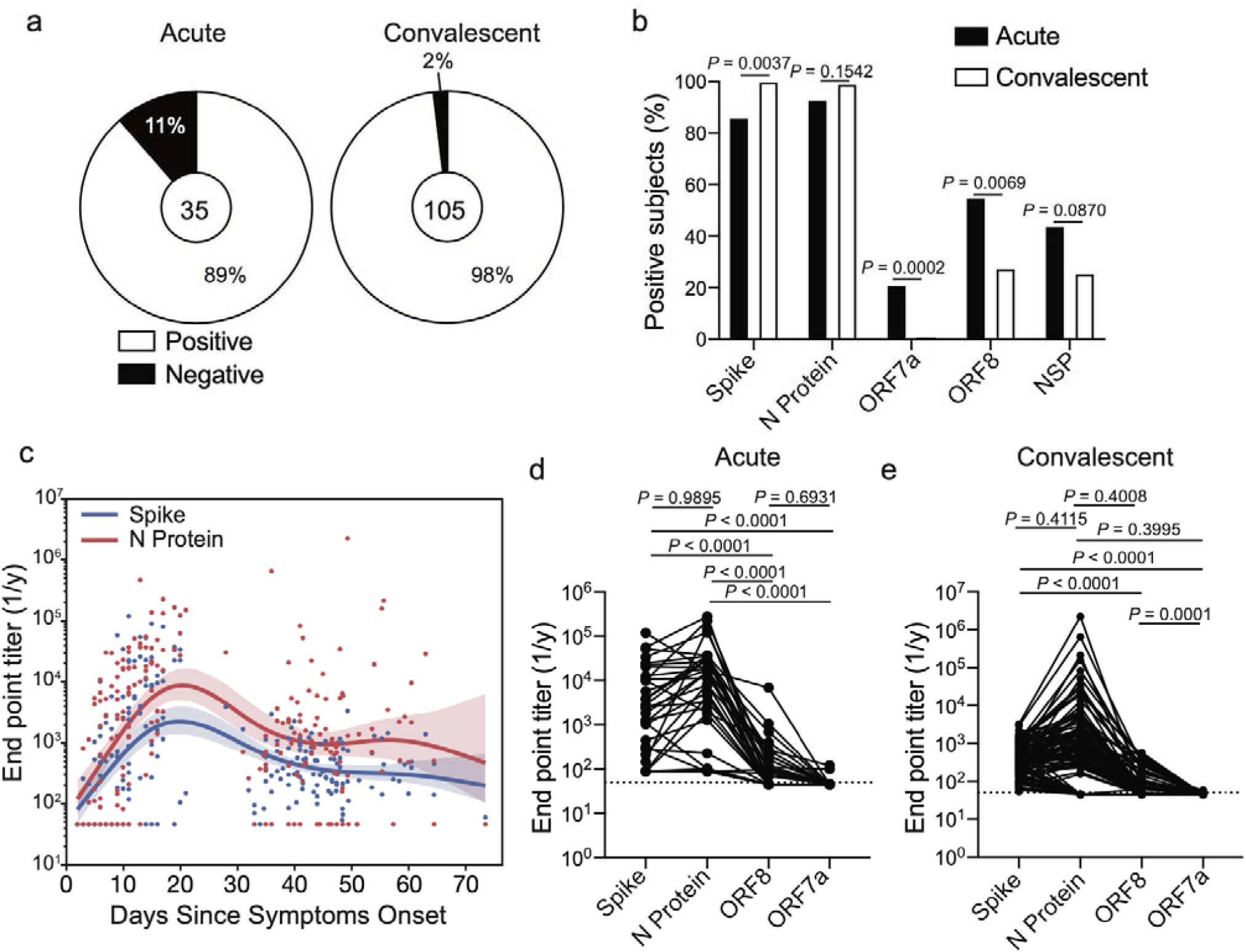
Antibody specificity and kinetics in SARS-CoV-2 infected subjects. **a**, Proportion of subjects in the acutely infected and convalescent cohorts who have seroconverted to one or more SARS-CoV-2 antigens. Number in center represents the number of subjects tested in each cohort. **b**, Proportion of subjects in the acutely infected (n=35) and convalescent (n=105) cohorts binding spike, N protein, ORF7a, ORF8, or at least one NSP antigen. **c**, Kinetics of plasma antibodies against the spike and N protein based on the start of symptoms. Data are pooled from the acute (n=117) and convalescent cohorts (n=105). Lines represent the fitted lines for spike and N protein titers and the shaded region indicates confidence of fit of the fitted line. **d** and **e**, end point titers of antibodies targeting spike, N protein, ORF7a, and ORF8 in the acutely infected cohort (**d**; n=35) and convalescent cohort (**e**; n=105). Lines connect titers across one subject. Data in **b** were analyzed using Fisher’s exact tests for statistical analyses. Data in **d** and **e** were analyzed using paired non-parametric Friedman tests. Dashed lines in **d** and **e** are the limit of detection.

To understand the inter-subject variability within our cohorts, we performed hierarchical clustering of subjects based on antibody titers against the spike, full length and RNA-binding domain of N protein, ORF7a, and ORF8 antigens. From the acutely infected cohort, we identified three clusters: high, mid, and low responders (Fig. 2a and Supplemental Table 3). Notably, the high responder cluster subjects were further from the onset of symptoms at the time of sampling and ultimately were hospitalized for a longer duration than those in the mid and low responder groups (Fig. 2b and c). We did not observe a statistical difference in age or sex between the three responder groups (Extended data Fig. 2a and b). Over 25% of subjects in the high responder group had a severe/highest CURB-65 score (Extended data Fig. 2c), a measure of pneumonia severity^9^, suggesting subjects in the high responder group had more severe infections. We further examined which features of the humoral immune response were driving subjects to segregate into these three clusters. Subjects within the high and mid responder groups robustly induced antibodies against the spike protein, but the high responder subjects mounted a larger response to N protein relative to the mid responder subjects (Fig. 2d-f). Additionally, subjects within the high and mid responder groups were more likely to mount an antibody response against ORF8 and NSP antigens (Extended data Fig. 2d and e). The low responder group largely did not mount an antibody response against any of the antigens tested (Fig. 2d-f and Extended Data Fig. 2d and e), although it is possible that plasma was collected before the subjects mounted a significant antibody response. Our data reveal that acutely infected subjects who were hospitalized for a longer duration mounted a larger antibody response against N protein and were more likely to mount a response against other SARS-CoV-2 antigens.

**Fig. 2:**
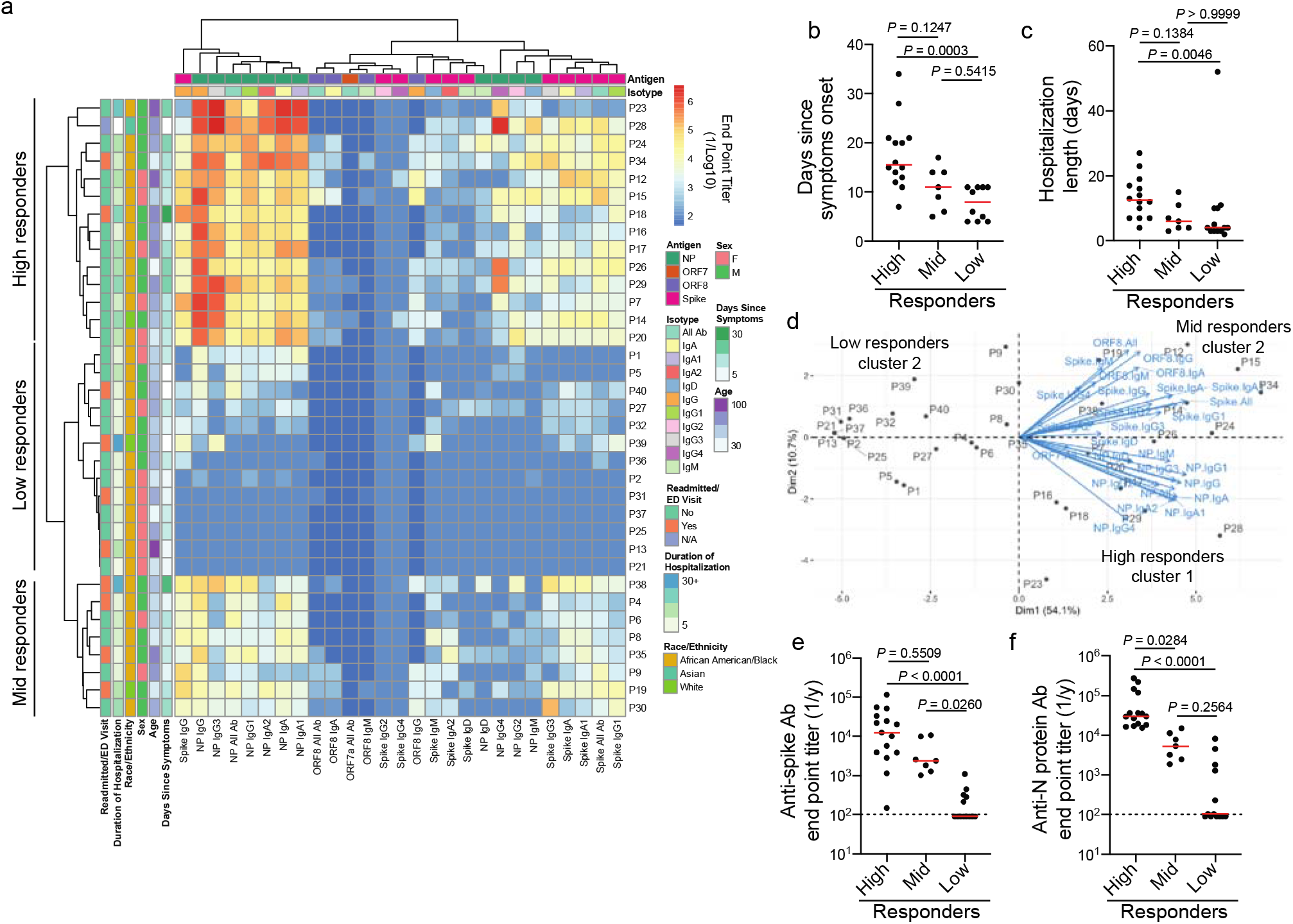
Acutely infected subjects with longer hospitalizations have a higher antibody response against N protein. **a**, Heatmap of hierarchical clustering of acutely infected subjects (n=35) based on antibody binding specificity and antibody isotype/subclass. Subjects clustered into three distinct clusters: high (n=15), mid (n=7), and low (n=13) responders. **b** and **c**, days since symptom onset (**b**) and length of hospitalization (**c**) amongst subjects in the high, mid and low responder clusters. **d**, PCA biplot of subjects clustering based on distinct antibody binding features. **e** and **f**, total antibody titers against the spike (**e**) and N protein (**f**) amongst the high, mid, and low responder clusters. Data in **b, c, e**, and **f** were analyzed using unpaired non-parametric Kruskal-Wallis tests. Dashed lines in **e** and **f** are the limit of detection. Bars in **b, c, e**, and **f** represent the median.

The convalescent cohort also clustered into three distinct clusters based on the magnitude of the antibody response against the spike and N protein (Fig. 3a and Supplemental Table 4), similar to the acutely infected cohort (Fig. 2). To understand the relationship between infection severity and antibody responses within the convalescent cohort, we scored subjects based on the severity and duration of self-reported symptoms and whether subjects were hospitalized (Supplemental Table 5). Notably, over 50% of subjects within the high responder group had a severe infection (Fig. 3b and Extended data Fig. 3a), indicating infection severity is linked to increased antibody titers. Moreover, subjects within the high responder group typically were older and male (Fig. 3c and d). Subjects within each responder group had a similar duration of symptoms (Extended data Fig. 3b), and subjects within all three groups had a similar amount of time to mount a response, as determined by the number of days since symptoms onset at the time of donation (Extended data Fig. 3c). Unlike the acutely infected cohort, subjects within the high responder group had higher titers against not only the N protein, but also the spike and ORF8 antigens relative to subjects within the mid and low responder groups (Fig. 3e and f, Extended data Fig. 3d and e), and were trending to be more likely to seroconvert against at least one of the NSP antigens tested (Extended data Fig. 3f). Consistent with these data, high and mid responder subjects had higher neutralizing titers than subjects in the low responder cohorts (Fig. 3g). In combination with the acutely infected cohort, our data reveal subjects with more severe infection are mounting a larger antibody response at both acute and convalescent time points.

**Fig. 3:**
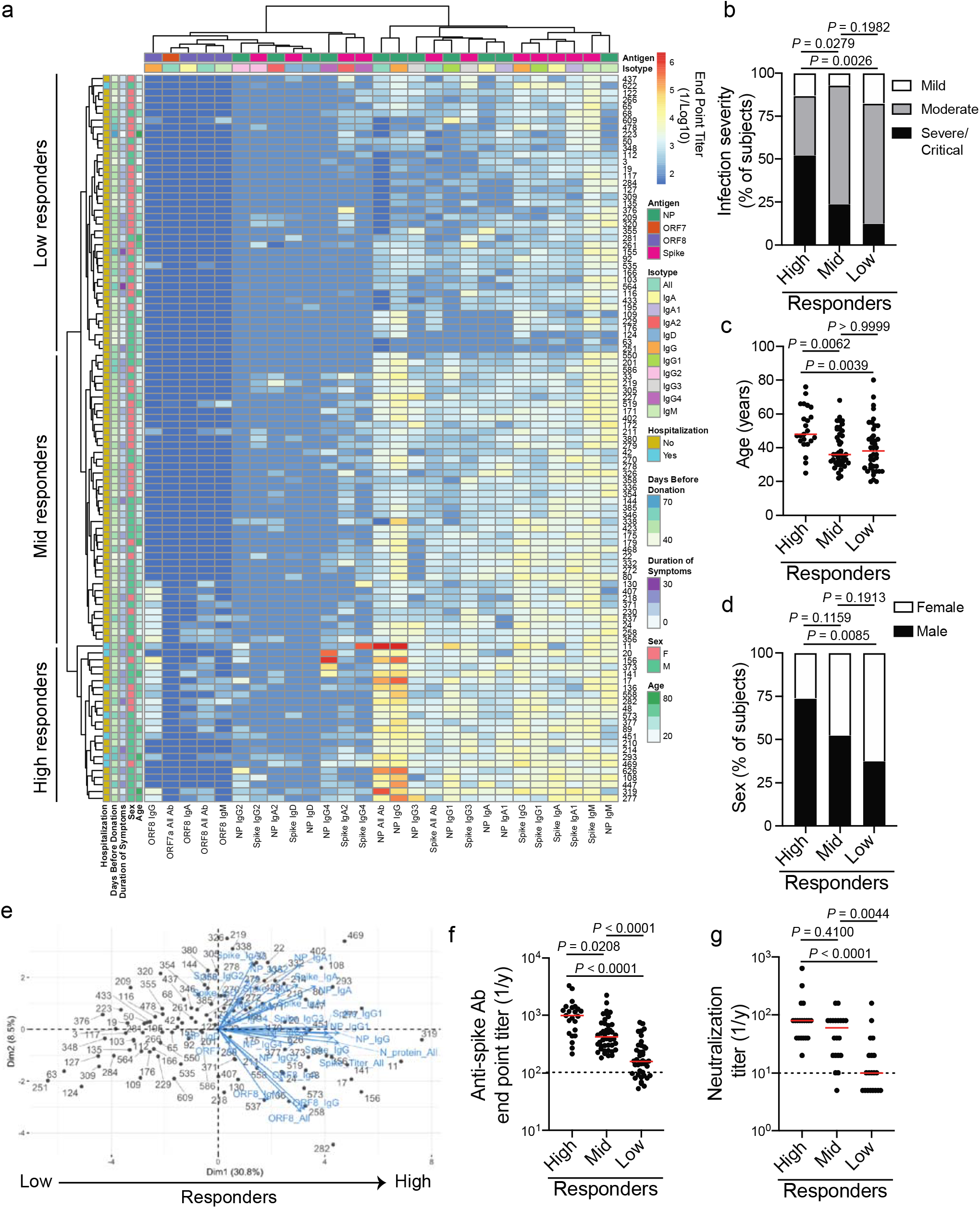
Convalescent subjects with higher antibody responses against multiple SARS-CoV-2 antigens tended to have a more severe infection. **a**, Heatmap of hierarchical clustering of convalescent subjects (n=105) based on antibody binding specificity and antibody isotype/subclass. Subjects clustered into three distinct clusters: high (n=23), mid (n=42), and low (n=40) responders. **b**-**d**, infection severity (**b**), age (**c**) and sex (**d**) of subjects in the high, mid and low responder clusters. **e**, PCA biplot of subjects clustering based on distinct antibody binding features. **f**, Total antibody titers against the spike amongst the high, mid, and low responder clusters. **g**, Neutralization titer, as determined by viral cytopathic effect, of 20 randomly selected samples from each of the high, mid, and low responder clusters. Data in **f** and **g** were analyzed using unpaired non-parametric Kruskal-Wallis tests. For **b-d**, data were analyzed using Fisher’s exact tests. Dashed lines in **f** and **g** are the limit of detection. Bars in **f** and **g** represent the median.

We next dissected the specificities of memory B cells (MBCs) induced by SARS-CoV-2 infection by performing B cell ELISpots on polyclonally stimulated peripheral blood mononuclear cells (PBMCs) isolated from convalescent subjects. Notably, MBCs largely targeted the spike, whereas very few MBCs targeted N protein or ORF8 (Fig. 4a). Additionally, subjects in the serum high responder group mounted a larger MBC response against the spike than subjects in the mid and low responder cohorts (Fig. 4b), with serum antibody titers against the spike positively correlating with the magnitude of the anti-spike MBC response (Fig. 4c). Despite the observed differences in anti-spike MBC responses between responder groups, we did not observe any differences in the anti-N protein and anti-ORF8 MBC response in the three responder cohorts (Extended data Fig. 4a and b). Together, these data indicate that the MBC response is largely directed against the spike protein, and that the high serum responder group mounted both a larger secreted antibody and MBC response upon SARS-CoV-2 infection.

**Fig. 4:**
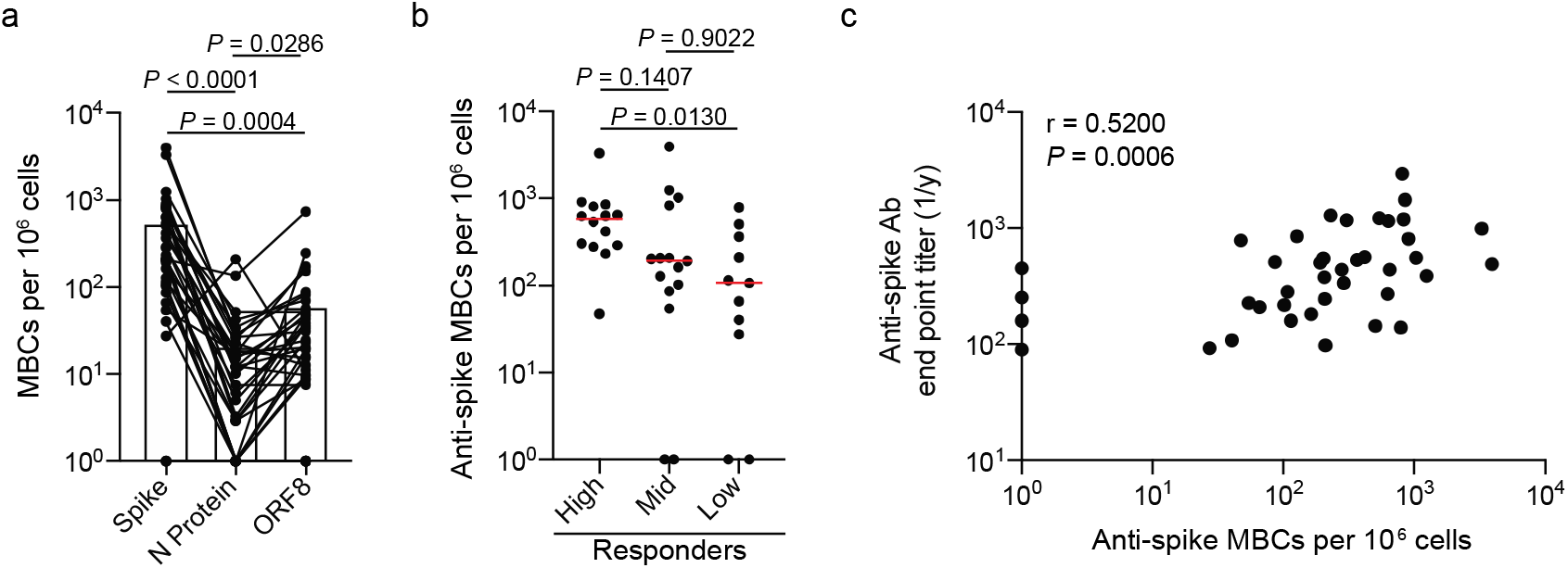
MBC response is largely driven against the spike. **a** and **b**, PBMCs from convalescent donors were polyclonally stimulated and ELISpots were performed to assess the number of antigen-specific MBCs. **a**, Number of MBCs (antigen-specific MBCs per 10^6^ cells) targeting the spike, N protein, or ORF8 (n=36). Lines connect antigen-specific MBCs across subjects. **b**, Number of spike targeting MBCs amongst the high (n=14), mid (n=15), and low responder (n=11) clusters. **c**, Spearman correlation of the number of anti-spike MBCs and anti-spike end point titers by individual (n=40). Data in **a** were analyzed using paired non-parametric Friedman tests. Data in **b** were analyzed using unpaired non-parametric Kruskal-Wallis tests. Data in **c** were analyzed by a non-parametric two-tailed Spearman correlation.

SARS-CoV-2 has acquired a D614G mutation within the spike protein and viruses carrying this mutation have since become the dominant circulating strain globally as of early April^10^. This mutation is located on the interface between two subunits of the spike trimer and may impact stability of the trimer^1^. As the subjects within our study were initially infected throughout March and into early April (Supplemental Tables 1 and 2), they were likely infected with the D614 variant. We did not observe a difference in antibody titers against the WT and D614G spike antigens within our acute cohort (Fig. 5a), suggesting the D614G epitope was not a major antigenic site. Strikingly, we identified that the convalescent cohort mounted a larger response against the G614 variant than the WT D614 that they were likely infected with (Fig. 5b), potentially due to the increased stability of the G614 variant^11^. Furthermore, we observed a strong positive correlation between D614 (WT) spike titers and G614 titers, indicating antibodies against the WT strain likely protect against the new G614 variant (Fig. 5c). These data indicate that the region that encompasses the D614G mutation is not immunodominant or does not affect the antigenicity of epitopes at or near this site. We also examined whether antibodies targeting the RBD of the spike protein cross-reacted with the RBD proteins of other pandemic threat coronaviruses, including SARS-CoV-1 and Middle East respiratory syndrome (MERS) CoV. We found a positive correlation between antibody titers against the SARS-CoV-2 RBD and the SARS-CoV-1 RBD, but not the MERS-CoV RBD (Fig. 5d and e). When divided by responder groups (Fig. 2 and 3), subjects in the high and mid responder groups had elevated titers against the SARS-CoV-1 RBD (Fig. 5f and g). These data show that subjects who mounted a larger response against the SARS-CoV-2 spike protein additionally mounted a larger antibody response against conserved epitopes that cross-react with closely related coronaviruses.

**Fig. 5:**
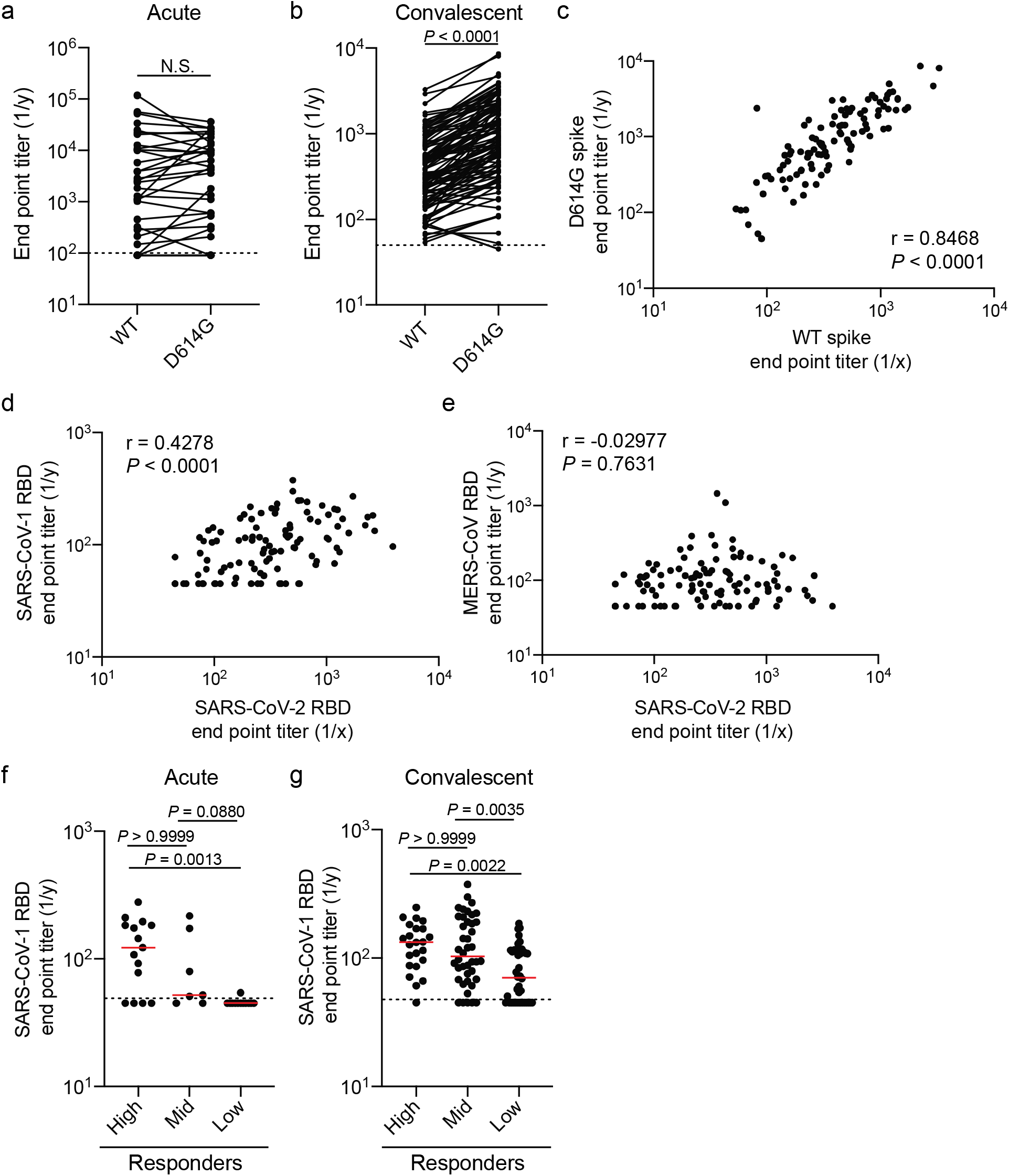
Antibody cross-reactivity to G614 spike mutant and SARS-CoV-1 and MERS-CoV RBD. **a** and **b**, End point titers of antibodies binding to the WT (D614) and mutant (D614G) SARS-CoV-2 spike protein from the acute (**a**; n=35) and convalescent (**b**; n=105) cohorts. **c**, Correlation of end point titers against the WT (D614) and mutant (D614G) spike from the convalescent cohort (n=105). **D** and **e**, correlation between SARS-CoV-2 RBD end point titers and SARS-CoV-1 RBD (**d**) or MERS-CoV RBD (**e**) end point titers from convalescent subjects (n=105). **f** and **g**, SARS-CoV-1 RBD end point titers amongst the high, mid, and low responder clusters from the acutely infected cohort (**f;** high n=23, mid n=42, and low n=40) and the convalescent cohort (**g**; high n=23, mid n=42, and low n=40). Data in **a** were analyzed using a two-tailed Wilcoxon matched-pairs signed rank test. Data in **b** were analyzed using a two-tailed paired t-test. For **c**-**e**, data were analyzed using a two-tailed Pearson correlation. Data in **f** and **g** were analyzed using unpaired non-parametric Kruskal-Wallis tests. Dashed lines in **a, b, f** and **g** are the limit of detection. Bars in **f** and **g** represent the median.

Together, our study demonstrates that severity of SARS-CoV-2 infection is associated with an increase in the magnitude and breadth of the ensuing humoral immune response. Notably, we identified the antibody response is largely mounted against the spike and N proteins, with the magnitude and kinetics of the anti-N protein antibody response outpacing the antibody response against spike. Although both proteins are highly expressed by coronaviruses, there is much more N protein as it encapsulates the whole viral genomic RNA, which is nearly 30 kb in size. As N protein dimer is projected to bind about 30 bp^12^, there are likely 1000+ N proteins per virion. In sharp contrast, there are only ∼26 spike trimers per virion^8^, suggesting the immunodominance towards N protein may be related to antigen burdens. Likewise, subjects with more severe disease likely have increased viral titers and free antigen in the lung lumen and draining lymph nodes, which could lead to increased antibody titers against nearly all antigens tested. Therefore, epitope spreading of the antibody response may be a factor of the amount of SARS-CoV-2 antigen present.

Subjects also mounted an antibody response against the accessory protein ORF8. ORF8 has immunoregulatory properties including the ability to limit type I interferon responses^13,14^ and downregulate MHC-I presentation to CD8 T cells^15^. Antibodies targeting ORF8 may limit these immunoregulatory properties, which could improve the host immune response and achieve better clinical disease outcomes. Additionally, we identified antibodies against non-structural proteins involved in viral replication, although antibodies against these antigens are unlikely to provide protection, as these antibodies targeting NSPs would need to be inside of a live cell while virus is replicating. Whether antibodies targeting discrete viral antigens other than the spike are neutralizing, have Fc-mediated effector functions, or are protective during infection remains to be determined.

Our study revealed acutely infected subjects who mounted higher antibody response relative to mid and low responder clusters tended to have higher pneumonia severity scores. Consistent with this notion, convalescent subjects who had higher antibody titers were those subjects who had a more severe infection. A recent report identified that subjects who succumbed to COVID-19 tended to mount a larger antibody response against N protein relative to the spike, whereas convalescent subjects tended to focus their antibody response on the spike protein^16^. However, our study identified that subjects generally had similar antibody responses against the N protein and spike, and infection severity was linked to an increase in antibody responses against both the spike and N protein. Ultimately, our findings on the relationship between infection severity and increased titers against the spike is consistent with a recent surveillance study performed in Iceland^6^.

The best clinical predictors of the magnitude of the antibody responses and epitope spreading within our convalescent cohort were age, sex, and hospitalization. Strikingly, the median age of the high responder cluster was 10+ years greater than the mid and low responder clusters (48 years vs. 36 and 38, respectively). Older adults are more likely to be symptomatic and hospitalized with SARS-CoV-2 infection^17,18^, suggesting increased disease severity and sustained viral titers over a longer period of time could lead to greater antibody titers against multiple viral antigens. Similarly, males were more likely to be segregated into the higher responder group despite the common finding that females generally mount higher antibody responses upon other viral infections and upon vaccination^19^. Although there is no difference in incidence of COVID-19 in men and women, men have a higher morbidity and mortality rate than women^20,21^ and likely experience increased viral titers and antigen persistence. Altogether, disease severity is the main clinical predictor of the magnitude of the antibody response mounted against SARS-CoV-2, as men and older adults are more likely to be hospitalized with COVID-19. It remains to be determined whether subjects with more severe disease are more likely to be protected from reinfection with SARS-CoV-2.

Together, our data indicate more severe infection is linked to a larger magnitude of circulating antibody and MBC response and increased viral antigen binding breadth across different viral antigens. CD4 T cells are critical for driving antibody responses by mediating germinal center selection of antigen specific B cells. Notably, CD4 T cells targeting multiple SARS-CoV-2 antigens and the magnitude of the CD4 T cell response positively correlate with SARS-CoV-2 specific antibody responses^22,23^. Moreover, subjects with more severe disease demonstrate an increased breadth and magnitude of the memory CD4 T cell response^23^, which could lead to the larger and broader antibody response of subjects with more severe infection, as observed in our study. The increase in the magnitude of the antibody response and MBC response in subjects with more severe infection could be due to increased CD4 T cell responses, although this was not directly tested in our study. However, subjects who succumbed to SARS-CoV-2 infection demonstrated a loss of germinal centers and CD4 T follicular helper cells^24^. These data in conjunction with our study suggest that an immunological balance will be needed to drive a sufficient secreted antibody response, MBC differentiation, and memory T cell responses that could provide robust protection from reinfection while preventing significant morbidity and mortality associated with SARS-CoV-2 infection.

## Methods

### Study cohorts

All studies were performed with the approval of the University of Chicago institutional review board and University of Chicago and University of Wisconsin-Madison institutional biosafety committees. Plasma samples from the acutely infected cohort were collected as residual samples submitted to the University of Chicago Medicine Clinical Laboratories. Convalescent subjects were recruited to donate one unit of blood for a convalescent plasma transfusion study, identified as clinical trial NCT04340050. 3 ml of blood and the leukoreduction filter were provided to the Wilson laboratory. All subjects in the acute and convalescent cohorts had PCR-confirmed SARS-CoV-2 infections.

### Recombinant proteins

Plasmids for the SARS-CoV-2 RBD and spike were provided by Dr. Florian Krammer at Icahn School of Medicine at Mount Sinai, and recombinant proteins were expressed in-house in HEK293F cells. D614G spike protein, SARS-CoV-1 RBD, and MERS-CoV RBD were generated in-house and expressed in HEK293F cells. ORF7a, ORF8, and full-length N proteins were cloned from the 2019-nCoV/USA-WA1/2020 SARS-CoV-2 strain at Washington University. Proteins were expressed in *Escherichia coli*, with N protein purified as a soluble protein and ORF7a and ORF8 oxidatively refolded from inclusion bodies. NSP antigens and the RNA-binding domain of N protein were provided by Dr. Andrzej Joachimiak at the Center for Structural Genomics of Infectious Diseases at the University of Chicago and Argonne National Laboratory and were expressed in *Escherichia coli*.

### Enzyme-linked immunosorbent assay (ELISA)

ELISAs performed in this study were adapted from previously established protocols^25,26^. Plasma samples were heat-inactivated for 1 hour at 56°C. High protein-binding microtiter plates (Costar) were coated with recombinant antigens at 2 μg/ml in phosphate-buffered saline (PBS) overnight at 4°C. Plates were washed with PBS 0.05% Tween and blocked with 200 μl PBS 0.1% Tween + 3% milk powder for 1 hour at room temperature. Plasma samples were serially diluted in PBS 0.1% Tween + 1% milk powder. Plates were incubated with serum dilutions for 2 hours at room temperature. Horseradish peroxidase (HRP)-conjugated goat anti-human Ig secondary antibody diluted in PBS 0.1% Tween + 1% milk powder was used to detect binding of antibodies, and after a 1-hour incubation, plates were developed with 100 μl SigmaFast OPD solution (Sigma-Aldrich), with development reaction stopped after 10 minutes using 50 μl 3M HCl. Absorbance was measured at 490 nm on a microplate spectrophotometer (BioRad). To detect binding of specific antibody isotypes and subclasses, ELISAs were performed using alternate secondary antibodies (Sigma-Aldrich; Jackson ImmunoResearch; Southern Biotech). End point titers were extrapolated from sigmoidal 4PL (where X is log concentration) standard curve for each sample. Limit of detection (LOD) is defined as the mean plus 2-8 S.D. (depending on antigen) of the O.D. signal recorded using plasma from SARS-CoV-2 negative human subjects. All calculations were performed in Prism 8 (GraphPad).

### Neutralization assays

Neutralization assays were performed by a viral cytopathic effect assay (CPE) using the SARS-CoV-2/UW-001/Human/2020/Wisconsin (UW-001), which was isolated from a mild human case in Wisconsin. Plasma was diluted 1:5 and serially diluted 2-fold and was mixed with an equal volume of virus (100 plaque-forming units) for a starting dilution of 1:10. The plasma/virus mixture was incubated for 30 minutes at 37°C and added to TMPRSS2-expressing Vero E6 cells grown in 1x minimum essential medium (MEM) supplemented with 5% FCS. Cells were incubated with plasma/virus mixture for 3 days, and then were fixed, stained, and analyzed. CPE was observed under an inverted microscope, and neutralization titers were determined as the highest serum dilution that completely prevented CPE.

### Memory B cell stimulations and enzyme-linked immunospot assays (ELISpot)

MBC stimulations were performed on peripheral blood mononuclear cells (PBMCs) collected from subjects in the convalescent cohort. To induce MBC differentiation into antibody secreting cells, 1×10^6^ PBMCs were stimulated with 10 ng/ml Lectin Pokeweed Mitogen (Sigma-Aldrich), 1/100,000 Protein A from *Staphylococcus aureus*, Cowan Strain (Sigma-Aldrich), and 6 μg/ml CpG (Invitrogen) in complete RPMI in an incubator at 37°C/5% CO_2_ for 5 days. After stimulation, cells were counted and added to ELISpot white polysterene plates (Thermo Fisher) coated with 4 μg/ml of SARS-CoV-2 spike that were blocked with 200 μl of complete RPMI. ELISpot plates were incubated with cells for 16 hours overnight in an incubator at 37°C/5% CO_2_. After the overnight incubation, plates were washed and incubated with anti-IgG-biotin and/or anti-IgA-biotin (Mabtech) for 2 hours at room temperature. After secondary antibody incubation, plates were washed and incubated with streptavidin-alkaline phosphatase (Southern Biotech) for 2 hours at room temperature. Plates were washed and developed with NBT/BCIP (Thermo Fisher Scientific) for 2-10 minutes, and reactions were stopped by washing plates with distilled water and allowed to dry overnight before counting. Images were captured with Immunocapture 6.4 software (Cellular Technology Ltd.), and spots were manually counted.

### Infection Severity Scoring and CURB-65 scoring

For the acutely infected cohort, CURB-65^9^ scores were calculated based on confusion, blood urea nitrate levels, respiratory rate, blood pressure, and age of subjects. For the convalescent cohort, we designed a severity scoring system (Supplemental Table 5) based on presence of 12 symptoms, duration of symptoms, and hospitalization, with a maximum of 35 points possible. Symptoms were scored based on presence or absence of 12 symptoms, severity (mild or moderate) of symptoms, with a possibility of 17 points. Duration of symptoms was broken down based on the number of weeks of symptoms. Hospitalized subjects were broken down based on oxygen supplementation and intensive care unit (ICU) admission. The criteria for scoring and the classification of certain scores (mild, moderate, severe, and critical infection) were determined before analyzing the data.

### Heatmaps, hierarchical clustering, and statistical analysis

Heatmaps were generated by ‘pheatmap’ *R* package (version 1.0.12). Features and subjects were clustered by the hierarchical clustering method implemented in the ‘pheatmap’ *R* package. Principal component analyses (PCA) were performed using ‘factoextra’ *R* package (version 1.0.7). Subjects were then visualized by their first two principal components (PC1 and PC2) on a 2D map. All statistical analysis was performed using Prism software (Graphpad Version 8), JMP (SAS Institute Version 15), or *R* (version 3.6.3). Specific tests for statistical significance used are indicated in the corresponding figure legends. *P* values less than or equal to 0.05 were considered statistically significant.

## Acknowledgments

This project was funded in part by the National Institute of Allergy and Infectious Diseases; National Institutes of Health grant numbers U19AI082724 (P.C.W.), U19AI109946 (P.C.W.), U19AI057266 (P.C.W.). This work was also partially supported by the National Institute of Allergy and Infectious Diseases Collaborative Influenza Vaccine Innovation Centers (CIVIC; 75N93019C00051, F.K. and P.C.W.), and the Centers of Excellence for Influenza Research and Surveillance (CEIRS) HHSN272201400008C (F.K.) and by the National Institute of Allergy and Infectious Diseases, National Institutes of Health, Department of Health and Human Services, under Contract HHSN272201700060C (A.J., D.F.). This work was also supported by the National Heart, Lung, Blood Institute award T32HL007605-35 (J.J.G.). We thank Shruti Kamath for assisting in analyzing CURB-65 scoring data for the acutely infected cohort. We would like to thank Dr. Robert Jedrzejczak for help with cloning, expression, and purification of SARS CoV-2 proteins. We are thankful to all subjects who participated in this study.

## Author Contributions

J.J.G. designed and performed experiments, analyzed the data, and wrote the manuscript. O.S. and J.W. performed experiments, analyzed the data, and wrote the manuscript. S.C., N.Y.Z., H.U., M.H., Y.N.D., C.A.N, and P.D.H. generated recombinant antigens. J.J.G., S.C., N.Y.Z., H.U., C.T.S., and H.L.D. processed convalescent blood samples. J.J.G., O.S., and C.T.S. processed samples from acutely infected subjects. J.J.G., J.W., S.C., C.T.S., and H.L.D. performed and analyzed ELISpot assays. L.L. performed hierarchical clustering and PCA analyses. P.H. and Y.K. performed neutralization assays. J.J.G., W.D.M., and M.L.M. designed the scoring system for convalescent subjects. M.J., K.S., J.S.D., and M.L.M. orchestrated convalescent plasma study. F.K., D.H.F., and A.J. provided recombinant antigens or plasmids to express recombinant antigens. V.T. provided samples from acutely infected subjects and helped analyze data. P.C.W. supervised the work. All authors edited the manuscript.

