## Supplemental Figures for "SARS-CoV-2 infection severity is linked to superior humoral immunity against the spike"

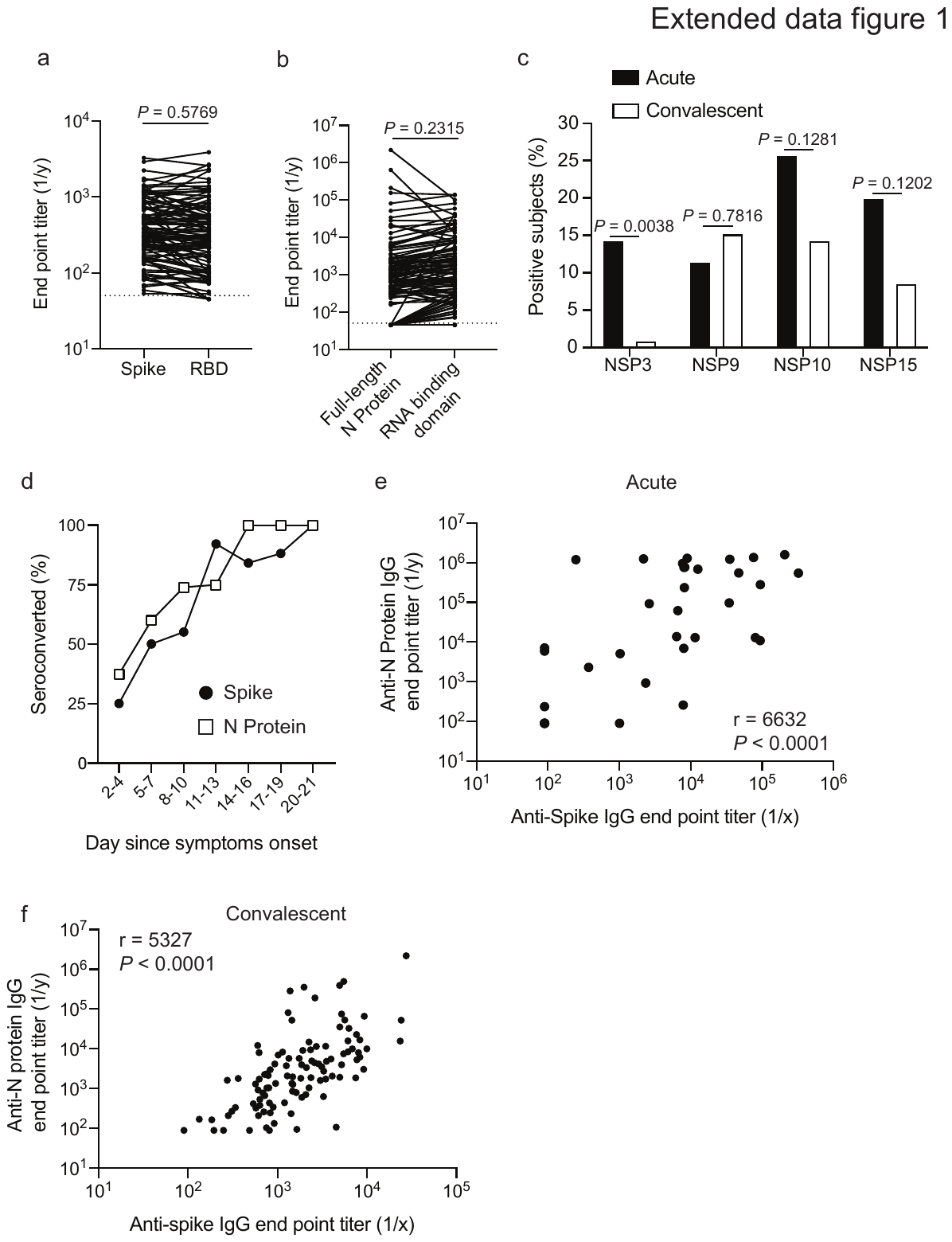

**Extended data Fig. 1: Specificity of serum antibody response of SARS-CoV-2 acutely infected and convalescent subjects. a** and **b**, End point titers of spike and RBD (**a**) and full-length N protein and RNA binding domain of N protein (**b**) from convalescent subjects (n=105). Lines connect titers from the same subject. **c**, Proportion of subjects with detectable antibodies against NSP antigens from the acute (n=35) and convalescent cohorts (n-105). **d**, Proportion of acutely infected subjects (n=35) who have seroconverted against the spike and N protein since days of symptom onset. **e** and **f**, correlation of anti-spike IgG and anti-N protein IgG titers in acute (**e**) and convalescent (**f**) cohorts. Data in **a** and **b** were analyzed using two-tailed paired t tests and data in **c** were analyzed using Fisher’s exact tests**.** Data in **e** were analyzed using a two-tailed Spearman correlation. Data in **f** were analyzed using a two-tailed Pearson correlation. Dashed lines in **a** and **b** are the limit of detection.

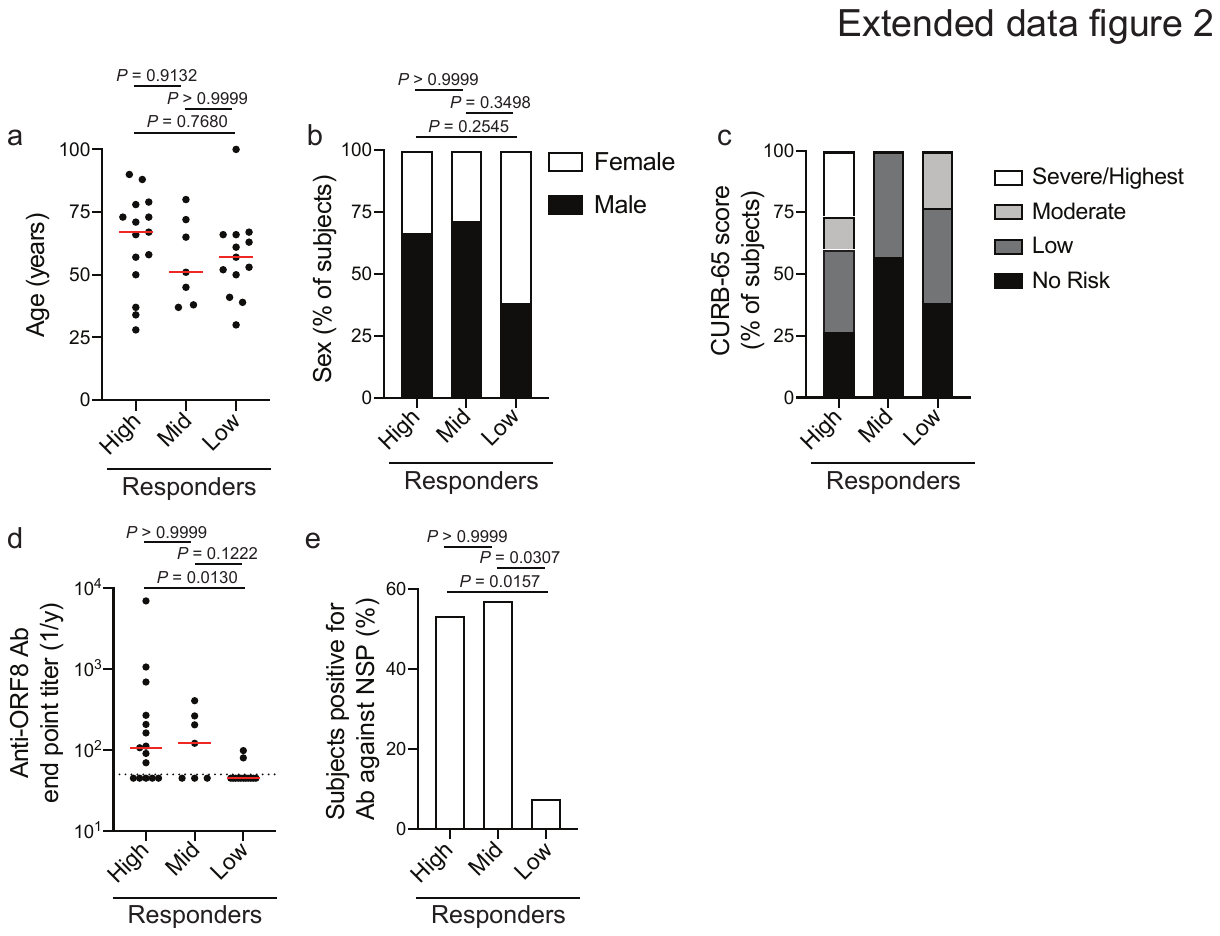

**Extended data Fig. 2: Clinical data and antibody specificity of acutely infected subject clusters. a**-**c**, Age (**a**), sex (**b**), and CURB-65 score (**c**) of subjects in the high (n=15), mid (n=7), and low (n=13) responder clusters. **d**, End point titers against ORF8 of subjects in the high (n=15), mid (n=7), and low (n=13) responder clusters. **e**, Proportion of subjects in the high (n=15), mid (n=7), and low (n=13) responder clusters with detectable antibodies against 1 or more NSP antigens. For **a** and **c**, data were analyzed by unpaired non-parametric Kruskal-Wallis tests. Data in **b** and **e** were analyzed by Fisher’s exact tests**.** Dashed lines in **d** are the limit of detection. Bars in **a** and **d** represent the median.

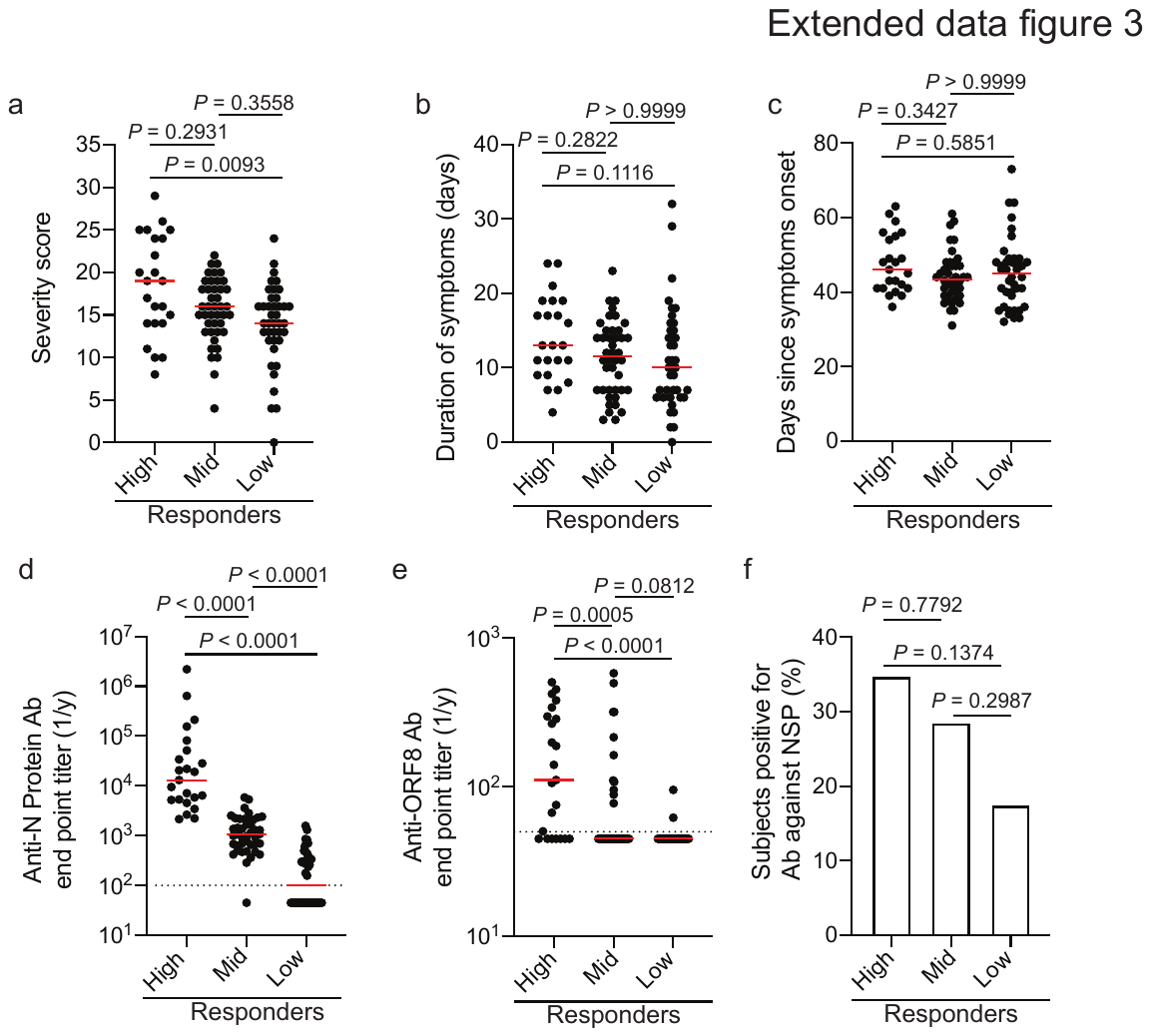

**Extended data Fig. 3: Clinical data and antibody specificity of convalescent subject clusters. a**-**c**, Severity score (**a**), duration of symptoms (**b**), and days since symptom onset (**c**) of subjects in the high (n=23), mid (n=42), and low (n=40) responder clusters. **d** and **e**, End point titers against N protein (**d**) and ORF8 (**e**) of subjects in the high (n=23), mid (n=42), and low (n=40) responder clusters. **f**, Proportion of subjects in the high (n=23), mid (n=42), and low (n=40) responder clusters with detectable antibodies against 1 or more NSP antigens. For **a**-**e**, data were analyzed by unpaired non-parametric Kruskal-Wallis tests. Data in **f** were analyzed by Fisher’s exact tests**.** Dashed lines in **d** and **e** are the limit of detection. Bars in **a**-**e** represent the median.

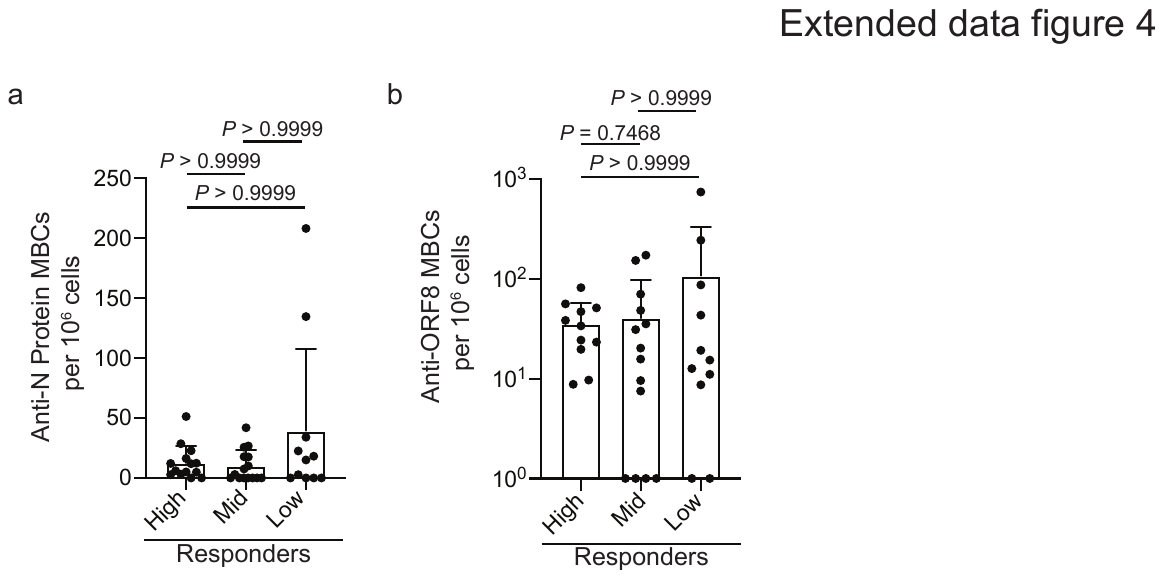

**Extended data Fig. 4: MBC responses against N protein and ORF8. a** and **b**, PBMCs from convalescent donors were polyclonally stimulated, and ELISPOTs were performed to assess the number of antigen-specific MBCs. **a**, Number of MBCs (antigen-specific MBCs per 10^6^ cells) targeting N protein (**a**) and ORF8 (**b**) amongst the high (n=14 for N protein; n=11 for ORF8), mid (n=16 for N protein; n=15 for ORF8), and low responder (n=10 for N protein and ORF8) clusters**.** For **a** and **b**, data were analyzed by unpaired non-parametric Kruskal-Wallis tests.

**Supplemental Table 1: Subject and clinical information for acutely infected cohort**

**Supplemental Table 2: Subject and clinical information for convalescent cohort**

**Supplemental Table 3**: ***P*-values between responder groups from the acutely infected cohort.** Related to Fig. 2. *P*-values of post-hoc pairwise comparisons of Analysis of Variance (ANOVA) between responder groups from the acutely infected cohort. *P* values were adjusted by Holm–Bonferroni method. *P* value = 0.00000 indicates *P* value < 0.00001. Red highlighted values represent statistically significant differences (*P ≤* 0.05) between groups.

|  | *P*-Values | | |  |
| --- | --- | --- | --- | --- |
| Antigen/Ab Isotype | High vs. Mid | Mid vs. Low | High vs. Low | Std. Deviation |
| NP IgG3 | 0.00000 | 0.00247 | 0.00000 | 1.65028627 |
| NP IgG | 0.00000 | 0.00000 | 0.00000 | 1.55238906 |
| NP IgA | 0.00000 | 0.00001 | 0.00000 | 1.39865608 |
| NP IgA1 | 0.00000 | 0.00010 | 0.00000 | 1.34647038 |
| NP IgA2 | 0.00000 | 0.00156 | 0.00000 | 1.25406493 |
| NP IgG1 | 0.00000 | 0.00000 | 0.00000 | 1.25071163 |
| NP IgG4 | 0.00067 | 0.63087 | 0.00003 | 1.24772617 |
| Spike IgG | 0.85674 | 0.00001 | 0.00000 | 1.12251522 |
| NP All Ab | 0.00270 | 0.00002 | 0.00000 | 1.11350163 |
| Spike All Ab | 0.03684 | 0.00004 | 0.00000 | 0.98688766 |
| Spike IgG3 | 0.53797 | 0.00046 | 0.00001 | 0.97972912 |
| Spike IgA | 0.28556 | 0.00266 | 0.00002 | 0.92272407 |
| ORF8 IgG | 0.41554 | 0.05648 | 0.00293 | 0.88124774 |
| Spike IgG1 | 0.04820 | 0.00023 | 0.00000 | 0.87884391 |
| NP IgM | 0.00002 | 0.19945 | 0.00000 | 0.87224077 |
| Spike IgA1 | 0.73044 | 0.00002 | 0.00000 | 0.86297411 |
| NP IgG2 | 0.00005 | 0.20110 | 0.00000 | 0.84540153 |
| Spike IgM | 0.22707 | 0.02034 | 0.11745 | 0.58655368 |
| NP IgD | 0.14321 | 0.37859 | 0.00964 | 0.5449614 |
| Spike IgA2 | 0.02472 | 0.00011 | 0.02472 | 0.51626759 |
| ORF8 All Ab | 0.37129 | 0.23313 | 0.01884 | 0.49777048 |
| ORF8 IgA | 0.39895 | 0.11318 | 0.00730 | 0.44250882 |
| Spike IgD | 0.86346 | 0.86346 | 0.37714 | 0.36506456 |
| Spike IgG4 | 0.41319 | 0.88793 | 0.32673 | 0.22016848 |
| ORF8 IgM | 0.94065 | 0.94065 | 0.48067 | 0.18630614 |
| ORF7a All Ab | 0.36390 | 1.00000 | 0.36390 | 0.09307027 |
| Spike IgG2 | 0.01960 | 0.45368 | 0.00092 | 0.01041245 |

**Supplemental Table 4**: ***P*-values between responder groups from the convalescent cohort.** Related to Figure 3. *P*-values of post-hoc pairwise comparisons of Analysis of Variance (ANOVA) between responder groups from the convalescent cohort. *P* values were adjusted by Holm–Bonferroni method. *P* value = 0.00000 indicates *P* value < 0.00001. Red highlighted values represent statistically significant differences (*P ≤* 0.05) between groups.

|  | *P*-Values | | |  |
| --- | --- | --- | --- | --- |
| Antigen/Ab Isotype | High vs. Mid | Mid vs. Low | High vs. Low | Std. Deviation |
| NP All Ab | 0.00000 | 0.00000 | 0.00000 | 0.97012491 |
| NP IgG | 0.00000 | 0.00000 | 0.00000 | 0.92822575 |
| NP IgG4 | 0.00006 | 0.29196 | 0.00000 | 0.65474305 |
| NP IgM | 0.44869 | 0.00063 | 0.00052 | 0.6466433 |
| ORF8 IgG | 0.00008 | 0.01025 | 0.00000 | 0.63588896 |
| NP IgG3 | 0.00211 | 0.00211 | 0.00000 | 0.63465636 |
| NP IgG1 | 0.00000 | 0.00000 | 0.00000 | 0.6337165 |
| Spike IgG3 | 0.00246 | 0.00246 | 0.00000 | 0.52020637 |
| Spike IgA1 | 0.46887 | 0.00000 | 0.00000 | 0.51098948 |
| Spike IgA | 0.52898 | 0.00000 | 0.00000 | 0.51020681 |
| Spike IgG1 | 0.00008 | 0.00001 | 0.00000 | 0.50629472 |
| NP IgA | 0.02180 | 0.00000 | 0.00000 | 0.50083625 |
| Spike IgG | 0.00027 | 0.00000 | 0.00000 | 0.49480188 |
| NP IgA1 | 0.03781 | 0.00000 | 0.00000 | 0.49265713 |
| Spike IgG4 | 0.35151 | 0.35151 | 0.11048 | 0.48422105 |
| Spike IgA2 | 0.72578 | 0.09653 | 0.28124 | 0.44303317 |
| Spike All Ab | 0.00021 | 0.00000 | 0.00000 | 0.39986788 |
| NP IgG2 | 0.00000 | 0.75031 | 0.00000 | 0.3439445 |
| Spike IgM | 0.98421 | 0.00668 | 0.01772 | 0.33589343 |
| ORF8 All Ab | 0.00024 | 0.01575 | 0.00000 | 0.31603601 |
| ORF8 IgM | 0.14538 | 0.14538 | 0.00870 | 0.28519883 |
| NP IgA2 | 0.73446 | 0.10013 | 0.10013 | 0.24871318 |
| Spike IgG2 | 0.70986 | 0.70986 | 0.38520 | 0.24461853 |
| ORF8 IgA | 0.00935 | 0.18456 | 0.00037 | 0.19149453 |
| Spike IgD | 0.80540 | 0.84585 | 0.80540 | 0.15863626 |
| NP IgD | 1.00000 | 1.00000 | 1.00000 | 0.05778103 |
| ORF7a All Ab | 0.81966 | 0.81966 | 1.00000 | 0.01082894 |

**Supplemental Table 5: Infection severity scoring system for convalescent subjects based on symptoms, hospitalization, and duration of symptoms.**
